# Engineering Antioxidant and Oxygen-Releasing Lignin Composites to Promote Wound Healing

**DOI:** 10.1101/2022.03.18.484913

**Authors:** Swathi Balaji, Walker D. Short, Benjamin W. Padon, Jorge A. Belgodere, Sarah E. Jimenez, Naresh T. Deoli, Anna C. Guidry, Justin C. Green, Tanuj J. Prajapati, Fayiz Farouk, Aditya Kaul, Dongwan Son, Olivia S. Jung, Carlos E. Astete, Myungwoong Kim, Jangwook P. Jung

## Abstract

The application of engineered biomaterials for wound healing has been pursued since the beginning of tissue engineering. Here, we attempt to apply functionalized lignin to confer antioxidation to the extracellular microenvironments of wounds and to deliver oxygen from the dissociation of calcium peroxide for enhanced vascularization and healing responses without eliciting inflammatory responses. Elemental analysis showed 17 times higher quantity of calcium in the oxygen releasing nanoparticles. Lignin composites including the oxygen releasing nanoparticles released around 500 ppm oxygen per day at least for 7 days. By modulating the concentration of the methacrylated gelatin, we were able to maintain the injectability of lignin composite precursors and the stiffness of lignin composites suitable for wound healing after photo-crosslinking. *In situ* formation of lignin composites with the oxygen releasing nanoparticles enhanced the rate of tissue granulation, the formation of blood vessels and the infiltration of α-smooth muscle actin^+^ fibroblasts into the wounds over 7 days. At 30 days after surgery, the lignin composite with oxygen generating nanoparticles remodeled the collagen architecture resembling to the reticular pattern of unwounded collagen with minimal scar formation. Thus, our study shows the potential of functionalized lignin for wound healing applications requiring balanced antioxidation and controlled release of oxygen for enhanced tissue granulation, vascularization and maturation of collagens.

## 1. INTRODUCTION

Wound healing is a critical process which progresses through tightly regulated phases and ultimately leads to repopulation of the wound with cells and extracellular matrix (ECM) to repair the injured site. A key aspect of the wound healing process involves the production of granulation tissue, a densely vascularized provisional tissue composed of fibroblasts (FBs), vascular endothelial cells (ECs), inflammatory cells and cell-derived ECM. Poor vascularization of the granulation tissue is often associated with impaired healing. Recent evidence further links excessive production of reactive oxygen species (ROS) and/or impaired detoxification of ROS to the pathogenesis of impaired wound healing[1–3]. Excess ROS accumulation disrupts cellular homeostasis and causes non-specific damage to critical cellular components and function, leading to impairment such as abhorrent FB collagen synthesis[4] and cell apoptosis, EC and smooth muscle cell dysfunction and compromised tissue perfusion, and increased proinflammatory cytokine secretion by macrophages[5]. Excess ROS is scavenged by enzymes such as superoxide dismutase and antioxidants that regulate the redox environment in healing skin wounds.

In recent years, antioxidants have drawn much attention as potential therapeutic interventions due to their ability to fight oxidative stress[6–12]. The main function of antioxidants is to scavenge or neutralize free radical formation and to inhibit the deleterious downstream effects of ROS. However, most antioxidants, taken orally, have limited absorption profile, which leads to low bioavailability and insufficient concentrations at the target site[13, 14]. To overcome this issue, research has been focused on developing tissue engineering strategies to provide locoregional delivery of antioxidants. Strategies including antioxidant nanoparticles made of inorganic materials such as mesoporous silica, cerium oxide, and fullerene, have been evaluated in *in vitro* assays and in animal models to determine their ability to scavenge free radicals while decreasing ROS concentrations to protect cells against oxidative stress[15–18]. Hydrogels that release ROS scavengers have been developed to promote cell function as a method to mitigate the foreign body response[19]. Additionally, decellularized myocardial matrix has shown to protect cardiomyocytes from ROS after myocardial infarction[20]. However, currently utilized biomaterials, especially natural biomaterials, employ relatively complex chemistry and lack mechanical tunability[21].

Lignin is a polyphenolic polymer that functions in plants to isolate pathogens to the site of infection while providing impermeability to cell walls[22]. Lignosulfonate is a water-soluble polymer derived from lignin and it decreases the nutritional quality of plant leaves for pathogens with structural damage, which eventually leads to the oxidation of lignosulfonate by hydrogen peroxide and inhibits the reproduction of bacteria[23–25]. We[26] and others[27] found that lignosulfonate can form nanostructures, which inspired us to apply lignosulfonate as a nanoscale carrier for drug or therapeutics while capitalizing on the inherent antioxidant properties of lignosulfonate. Successful application of engineered biomaterials for wound healing relies upon overcoming the limitation of oxygen diffusion via exogenous oxygenation such as using perfusion bioreactors. In a small scale oxygenation, many solid inorganic peroxides have been used to support cell growth, survival, tissue regeneration, and bioremediation[28]. Calcium peroxide (CaO_2_) possesses many distinctive properties in comparison with other peroxides, including better thermal stability, environmental harmless end products, extended release period of hydrogen peroxide and reasonable cost[29, 30]. Calcium and magnesium peroxide have low solubility coefficients, 1.65 mg/mL at 20°C and 0.86 mg/mL at 18°C, respectively[31]. Based on these values, calcium peroxide has a higher oxygen-generation potential than magnesium peroxide[31, 32].

Therefore, to apply antioxidation and locoregional oxygenation, we developed injectable lignin composites with the following components and properties: 1) thiolated lignosulfonate (TLS) scavenges ROS from wounds[33, 34], 2) methacrylated gelatin (GelMA) modulates the mechanical properties[34] of lignin composites to support injectability, and 3) sodium lignosulfonate (SLS)[26] encapsulates CaO_2_ while scavenging radicals from CaO_2_[26, 35] and simultaneously protecting CaO_2_ from aqueous environment of tissue or hydrogel. First, we assessed the integration of CaO_2_ to SLS-PLGA (poly(lactic-*co*-glycolic) acid) nanoparticles (NPs) and the release of O_2_ from lignin composites. Then, swelling/degradation profiles and mechanical properties of lignin composites were assessed. Lastly, we applied lignin composites in the wounds of wildtype mice and assessed tissue granulation, neovascularization and wound healing process, inflammatory responses and scaring.

## 2. MATERIALS AND METHODS

### 2.1 Synthesis of NPs

The coupling of lignosulfonate to PLGA was performed by acylation reaction for SLS[26] in a mass ratio of 2 to 1. The final precipitate was collected and dried for 2 days under vacuum. Then, SLS-*graft*-PLGA (SLS-PLGA) was dissolved in ethyl acetate (EtOAc, organic phase) and calcium peroxide (CaO_2_, Cat# 466271, Sigma-Aldrich, St. Louis, MO, USA) and dissolved under stirring at room temperature. The organic phase was added to the aqueous phase (low resistivity water) under stirring. The suspension was homogenized with a microfluidizer (Microfluidics Corp., Westwood, MA, USA) at 30 kpsi at 4°C. Next, the organic solvent was evaporated with a rotary evaporator R-300 (Buchi Corp, New Castle, DE, USA). Finally, trehalose was added (1:1 mass ratio) as a cryoprotectant for the lyophilization of SLS-PLGA NPs and stored at −20°C for further characterization and experiments.

### 2.2 Dynamic light scattering (DLS) of NPs

Size, polydispersity and ζ-potential of NPs of CPO and TLS (0.2-0.4 mg/mL) were measured by dynamic light scattering (DLS) using Malvern Zetasizer ZS (Malvern Panalytical, Westborough, MA, USA). NP suspensions were filtered through 0.45 μm filter. To prevent the formation of disulfide bonds, TLS sample was prepared in 250 mM Tris(2-carboxyethyl)phosphine hydrochloride (TCEP HCl, Cat# H51864, Alfa Aesar, Haverhill, MA, USA) in PBS.

### 2.3 Elemental analysis of NPs

We utilized the nuclear microscopy setup at the Louisiana Accelerator Center to probe the concentrations of calcium in our samples using particle-induced x-ray emission (PIXE) spectrometry[36]. The samples were placed in a low-pressure environment (≤1×10^−6^ mbar) on electrically conductive non-porous carbon tape attached to the sample holder. A focused (10 μm 10 μm) 2 MeV proton beam, with a beam current in the range of 10−20 pA, raster scanned the sample region (1 mm × 1 mm) for about 1 h. A silicon drift detector was placed at 135° in front of the sample to detect the characteristic x-rays excited by energetic protons. Analysis of the spectra was performed with the GeoPIXE™ software (v7.3)[37]. The elemental maps and the derived concentrations were generated by the dynamic analysis method using average matrix composition from the whole scanned area.

### 2.4 Formation of lignin composites

TLS was synthesized with the previously published protocol[33]. GelMA was synthesized by the coupling reaction of gelatin with methacrylic acid (see supplementary information for details). We confirmed the alkene incorporation to gelatin with ^1^H nuclear magnetic resonance (NMR, **Figure S1**). Lignin composites were formed by weighing out GelMA, lithium phenyl-2,4,6-trimethylbenzoylphosphinate (LAP, Allevi, Philadelphia, PA, USA) and NPs of TLS, SLS-PLGA/CaO_2_ or SLS-PLGA (w/o CaO_2_) as summarized in **Table 1**. Any concentrations of TLS higher than 7 mg/mL interfered photo-crosslinking of lignin composites, leading to the reduction of elasticity of lignin composites with partially crosslinked lignin composites. The concentrations of GelMA, LAP and TLS were fixed at 50, 5 and 3 mg/mL, respectively and the concentrations of SLS-PLGA/CaO_2_ or SLS-PLGA (w/o CaO_2_) were varied at either 4 or 40 mg/mL. The precursor of lignin composites was prepared by mixing GelMA with LAP and NPs in PBS at 37°C as summarized in **Table 1**. Each precursor was pipetted into a custom polydimethylsiloxane (PDMS) mold (8 mm diameter and 1 mm height, 50 μL) to form lignin composites. Samples were crosslinked using an UV floodlamp (Intelli-Ray 400, Uvitron international, West Springfield, MA, USA) for 30 s at 10 mW/cm^2^. To prevent confusion, abbreviation of lignin composites is summarized in **Table 1**.

**Table 1.** Abbreviation of lignin composites

| Abbreviation | Matrix | Nanoparticles (NPs) |
| --- | --- | --- |
| UNTX | None | None |
| GelMA | GelMA | None |
| TLS | GelMA | TLS |
| CPOc | GelMA | TLS and SLS-PLGA (w/o CaO<sub>2</sub>) |
| CPO | GelMA | TLS and SLS-PLGA/CaO<sub>2</sub> |

### 2.5 Quantification of O_2_ release from lignin composites

The released O_2_ was optically measured by a planar O_2_ sensor spot (SP-PSt3-SA23-D5-OIW-US, PreSens, Regensburg, Germany) placed at the bottom of a 96-well plate. Lignin composites (5 mm diameter and 1 mm height, 20 μL) were placed on top of the planar O_2_ sensor spot and submerged in serum-free medium (37°C/5% CO_2_). To prevent evaporation, each well was covered with parafilm and the entire plate was wrapped with parafilm. A polymer optical fiber (POF-L2.5-2SMA, PreSens, Regensburg, Germany) optically sensed the concentration of O_2_ through the bottom of a well and sent signals to an O_2_ sensor (OXY-1 SMA, PreSens, Regensburg, Germany)[38] for measurement.

### 2.6 Swelling ratio and *in vitro* degradation of lignin composites

Swelling ratios of lignin composites were determined by the following equation: [W_s_-W_d_]/W_d_, where W_s_ and W_d_ represent the weight after swelling in PBS for 24 h and the weight after lyophilization, respectively. *In vitro* degradation of lignin composites was determined by submerging lignin composites in the solution of collagenase type II (0.5 U/mL, cat# CLS-2, Worthington Biochemical) with 1 mM CaCl_2_ in serum-free culture medium, along with a control group without including the collagenase. Lignin composites were collected at 0, 2, 6 and 24 h and lyophilized to determine the fraction of remaining composites.

### 2.7 Oscillating rheometry of lignin composites

Using a TA Discovery HR-2 rheometer and the previously published methods[33], viscosity of the precursors of lignin composites were measured in a flow ramp setting (shear rate from 1 to 2000 (1/s)) and with a 25 mm parallel plate. Using an 8 mm parallel plate, storage (G′) and loss (G″) moduli of lignin composites were determined by frequency sweeping from 0.62 to 19.9 (rad/s) at 2% strain. Since storage moduli are altered by axial stress applied during measurement, we evaluated the slope of axial stress vs compression, similar to evaluating Young’s modulus from the slope of a stress-strain curve[39]. Axial stresses at 0, 10 and 20% of compression were determined, while lignin composites were subject to 2% strain and 6.28 rad/s frequency.

### 2.8 Animal model of wound healing

Wound healing studies were carried out in wildtype (WT) C57BL/6N mice (8- to 10-week-old, female and male). Mice were maintained under pathogen-free conditions with access to food and water ad libitum in the Texas Children’s Hospital Feigin Center animal facility. Protocols for animal use were approved by the Institutional Animal Care and Use Committee at Baylor College of Medicine. At the time of wounding, mice were anesthetized using isoflurane, the backs were shaved, and prepped for surgery with three times alternating betadine and 70% isopropyl alcohol scrubbing. Two 6 mm diameter full thickness wounds were made using a 6 mm dermal punch biopsy, excising tissue through the panniculus carnosus muscle. Immediate skin contraction was controlled through application of a silicone stent with inner diameter of 8 mm and outer diameter of 16 mm, secured concentric to wound using skin adhesive and 6 simple interrupted 6-0 Proline© sutures (Ethicon, Raritan, NJ). Wounds were maintained in a moist wound environment using a semi-occlusive sterile adhesive dressing (Tegaderm, 3M, St. Paul, MN, USA) and contraction was controlled through application of a silicone stent with inner diameter of 8 mm and outer diameter of 16 mm, placed concentric to wound using skin adhesive and 6 interrupted sutures. Prior to dressing with Tegaderm, each wound received one of four treatments: a standard saline wash (untreated, UNTX), TLS, CPOc and CPO (summarized in **Table 1**). Crosslinking of the treatments was performed immediately after application to the wound bed using a UV floodlamp (B-100AP High Intensity, Blak-Ray) for an exposure time of 30 s. Wounds were imaged at 1, 3, and 7 days post operatively (**Figure S2**), and harvested at days 7 and 30. For histology and immunostaining, wounds were bisected in the rostral-caudal plane, fixed overnight in 10% neutral buffered formalin, dehydrated through a series of graded ethanol and xylene, and embedded in paraffin wax. Five-micrometer-thick sections from the paraffin-embedded wounds were collected using a RM 2155 microtome (Leica, Heidelberg, Germany) and used in staining.

### 2.9 Morphometric quantification

At day 7, epithelial gap and granulation tissue areas were measured from Hematoxylin and Eosin (Cat#3801571 and Cat#3801615, respectively; Leica, Heidelberg, Germany) stained sections using morphometric image analysis (LASX, Leica, Heidelberg, Germany). Staining was carried out on 5 μm formalin fixed paraffin embedded (FFPE) sections following deparaffinization and rehydration in graded xylene and ethanol as following the manufacturer’s recommendations. Epithelial gap was determined using a full, 4× tile scan of the wound bed, and measuring the distance (in mm) between the leading epithelial margins on either side of the wound. Granulation tissue area was determined using a standardized approach of calculating the entire wound area (in mm^2^) above the panniculus carnosus and bounded within the wound edges laterally. If parts of the hydrogel remained devoid of infiltrated cells, that area was excluded from the granulation tissue assessment.

### 2.10 Immunostaining

Immunohistochemistry was carried out on serial 5 μm sections from day 7 FFPE tissues, which were deparaffinized and rehydrated to water using graded xylene and ethanol. Primary antibodies used were αSMA (rabbit anti-mouse, Cat# ab5694, Abcam, Cambridge, United Kingdom, 1:500 dilution) to measure myofibroblast infiltration, CD31 (rabbit anti-mouse, Cat# ab28364, Abcam, Cambridge, United Kingdom, 1:100 dilution) to measure endothelial cells and vascular lumen density, CD45 (rabbit anti-mouse, Cat# ab10558, Abcam, Cambridge, United Kingdom, 1:5000 dilution) to determine pan-leukocyte infiltration, CD206 (rabbit anti-mouse, Cat# ab64693, Abcam, Cambridge, United Kingdom, 1:500 dilution) to measure M2 macrophage levels, F4/80 (rat anti-mouse, Cat# MF48000, Thermo Fisher Scientific, Waltham, MA, USA, 1:5000 dilution) to measure pan macrophage infiltration, and Ly6G (rat anti-mouse, Cat# 551459, BD Biosciences, Franklin Lakes, NJ, USA, 1:5000 dilution) to determine infiltration of monocytes, granulocytes, and neutrophils.

Following deparrafinization and rehydration to water, sections were then immersed in target antigen retrieval solution (Cat #K800521-2, Agilent, Santa Clara, CA, USA) and treated following the protocol within the DAKO PT Link Rinse Station (Cat #PT10930, Leica, Heidelberg, Germany). Following antigen retrieval, wound sections were stained using the DAKO autostainer (AS480, Leica, Heidelberg, Germany). First, the sections were buffered in DAKO wash buffer (Cat #K800721-2, Agilent, Santa Clara, CA, USA) before incubating in primary antibody diluted in DAKO antibody diluent (Cat# S080983-2, Agilent, Santa Clara, CA, USA) for 1 h. The primary antibody was then rinsed, and the sections were washed in wash buffer before incubation in the appropriate the secondary antibody system, either HRP-Rabbit (Cat# K400311-2, Agilent, Santa Clara, CA, USA) or HRP-Rat (Cat#D35-110, GBI Labs, Bothell, WA) for 20 min. The secondary antibody was then rinsed off using wash buffer, and the appropriate visualization system was applied – DAKO DAB (Cat# K346811-2, Leica, Heidelberg, Germany) for all but CD206 which utilized AEC (Cat# K400511-2, Leica, Heidelberg, Germany). A hematoxylin (Cat #K800821-2, Agilent, Santa Clara, CA, USA) counterstain was then applied, and sections were dehydrated in graded xylene and ethanol before putting coverslips using a xylene based mounting media (Cat#23-245691, Thermo Fisher Scientific, Waltham, MA, USA) – except for those sections using AEC as the visualization system which required an aqueous mounting media (Cat#108562, Merck KGaA, Darmstadt, Germany).

Staining was quantified using images taken on the Leica DMI8 camera. For all data, the percentage of positive cells were determined by counting the number of positive cells per High Powered Field (HPF, 40× magnification) and dividing by the total number of cells in that HPF as determined by the hematoxylin counterstain. Final values were determined by the average of 4 to 6 HPFs per wound. Total number of vessel lumens were counted per HPF. These percent values or vessel counts were then averaged from 6 images taken across the wound bed to determine the final value.

### 2.11 Scar assessment

Serial 5 μm sections from D30 FFPE tissue were deparaffinized and rehydrated to water using graded xylene and ethanol. Sections were then stained using Gomori’s Trichrome (blue) as following the manufacturer’s recommendations (Cat# 38016SS2, Leica, Heidelberg, Germany). Collagen content per HPF in the dermis of the scars was measured using established methods with color-thresholding in ImageJ in which color segmentation was used to isolate only blue pixels, representing collagen fibers, thus allowing quantification of the amount of collagen within the selected area[40]. Gross images of the wound at day 30 were also obtained, and a subjective assessment of scarring was performed.

### 2.12 Statistical analysis

For multiple comparisons, one-way ANOVA (analysis of variance) with Tukey’s *post hoc* tests or with Kruskal-Wallis test followed by Dunn’s test was performed, where p values <0.05 or <0.01 were considered significant. At least three independent experiments were performed.

## 3. RESULTS

### 3.1 The incorporation of CaO_2_ in NPs of SLS-PLGA led a narrower distribution of NP diameters

Our effort to apply the antioxidant capability of TLS to wound microenvironments would be synergistic with other pro-regenerative stimuli. We hypothesized that controlled release of oxygen while scavenging ROS by TLS in the wound microenvironments will promote wound healing. Thus, we incorporated CaO_2_ in NPs of SLS-PLGA/CaO_2_ [26] and found that the size of NPs is not significantly altered, even when compared to NPs of SLS-PLGA (w/o CaO_2_), as shown in **Table 1**. In comparison to SLS, the average diameter of NPs of SLS-PLGA/CaO_2_ increased while the PdI of NPs of SLS-PLGA/CaO_2_ was significantly smaller than that of NPs of SLS-PLGA (w/o CaO_2_). SLS is natural biomaterial with potential lot-to-lot variability. However, the synthesis of NP with CaO_2_ slightly increased the average diameter and significantly reduced PdI upon the incorporation of CaO_2_. In contrast, NPs of SLS-PLGA (w/o CaO_2_) were formed only with ethyl acetate, forming irregular shapes possibly with cavity. The average diameter of TLS is significantly smaller than SLS or NPs of SLS-PLGA. After adding TCEP to cleave possible disulfide bonds in TLS, the average diameter of TLS was further reduced. As evidenced in **Table 2**, TLS formed NPs with a broader distribution. The formation of NPs of SLS-PLGA requires stirring at room temperature, while the thiolation of SLS is completed via acid-catalyzed esterification at 80°C. In PBS (pH 7.4), NPs were formed via interparticle disulfide formation with thiols in TLS, but adding TCEP to PBS dissociates NPs of TLS into smaller NPs. This could be an advantage to form lignin composites with homogeneous distribution of TLS in the matrix of GelMA. After the degradation of GelMA, reduced form of TLS can be cleared by glomerular filtration (5-10 nm).

**Table 2.**
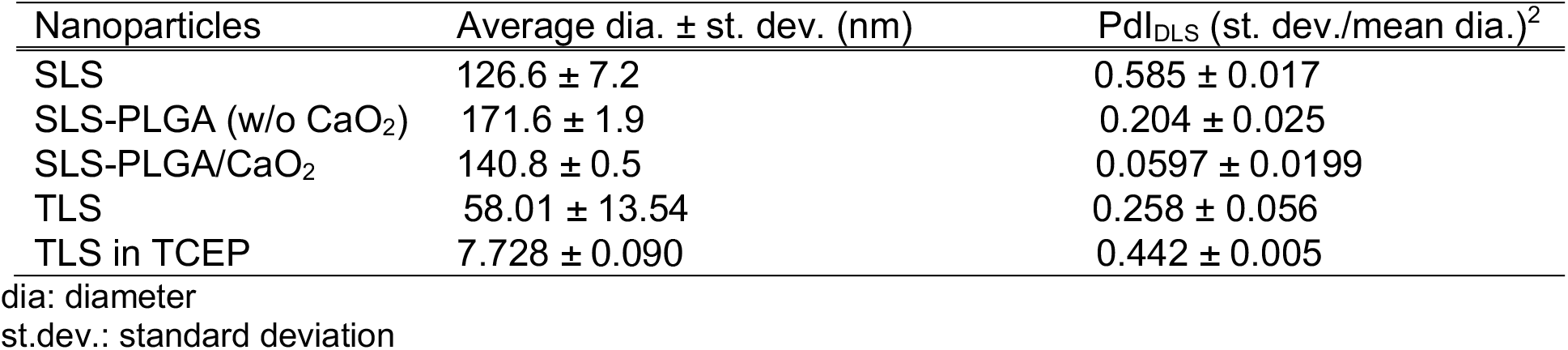
DLS of NPs.

| Nanoparticles | Average dia. ± st. dev. (nm) | Pdl<sub>DLS</sub> (st. dev./mean dia.)<sup>2</sup> |
| --- | --- | --- |
| SLS | 126.6 ± 7.2 | 0.585 ± 0.017 |
| SLS-PLGA (w/o CaO<sub>2</sub>) | 171.6 ± 1.9 | 0.204 ± 0.025 |
| SLS-PLGA/CaO<sub>2</sub> | 140.8 ± 0.5 | 0.0597 ± 0.0199 |
| TLS | 58.01 ± 13.54 | 0.258 ± 0.056 |
| TLS in TCEP | 7.728 ± 0.090 | 0.442 ± 0.005 |

dia: diameter
st.dev.: standard deviation

Due to the size (diameter) of NPs, we assessed the incorporation of CaO_2_ by PIXE spectrometry. NPs of SLS-PLGA with or without CaO_2_ showed similar composition of elements while the normalized concentration of Ca in NPs of SLS-PLGA/CaO_2_ is about 17 times higher than that of NPs of SLS-PLGA (w/o CaO_2_) in the quantitative molecular spectroscopy (**Figure 1**).

**Figure 1.**
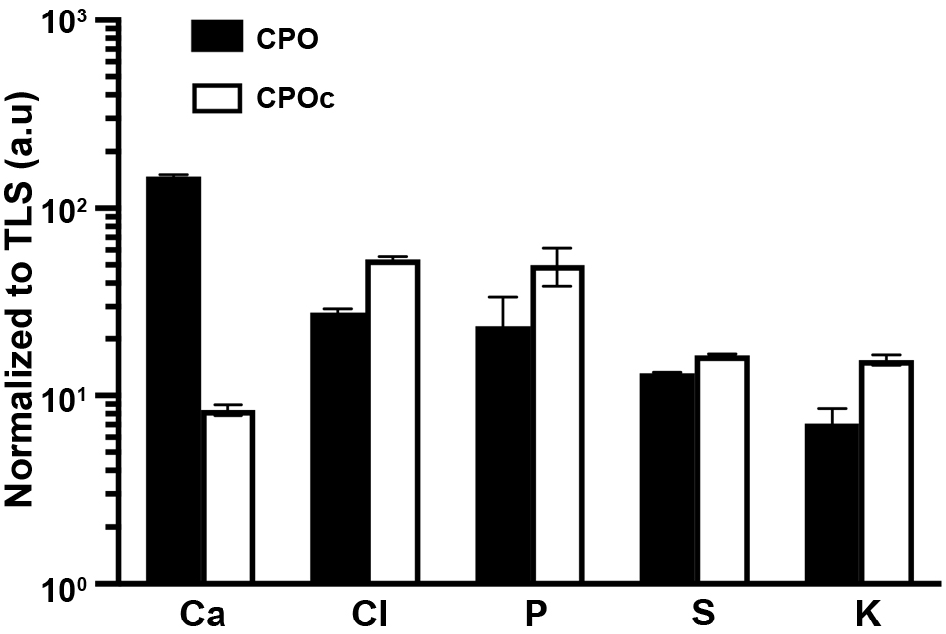
Elemental analysis of NPs of SLS-PLGA with or without CaO_2_ by microPXIE. The concentration (arbitrary units, a.u.) of Ca, Cl, P, S and K in the NPs normalized to the NPs of TLS. TLS also had trace amounts of Mn and Fe. The bars represent counting uncertainties (1σ) in a single measurement analyzed by GeoPIXE software; mean±SEM.

### 3.2 Oxygen released from CPO lignin composites was maintained at around 700 ppm/day from each composite for 7 days

To measure the released quantity of O_2_ from lignin composites, we used a planar O_2_ sensor with optical measurement. These planar sensors can measure the concentration of O_2_ in liquid or gas. The difference (Δ) of the area under curve (AUC) between lignin composites with NPs of SLS-PLGA/CaO_2_ and of SLS-PLGA (w/o CaO_2_) was calculated over 1440 min each day. As shown in **Figure 2**, the difference is around 500 ppm (0.05% O_2_) per day from the lignin composite with 5 mm diameter and 1 mm height (20 μL). This amount can be scaled to 700 ppm per day with lignin composites (6 mm diameter and 1 mm height) for animal experiments. As the statistical difference is not detected, the oxygen release is maintained up to day 7, which is also distinguished from other methods[41–46] of O_2_ delivery by CaO_2_. We observed that the swelling of lignin composites over the first 24 h contributed to the slightly higher ΔAUC in day 1 than that in day 2 since lignin composites were placed in a well of the 96-well plate with the planar O_2_ sensor and serum-free medium was added without achieving equilibrium swelling of lignin composites.

**Figure 2.**
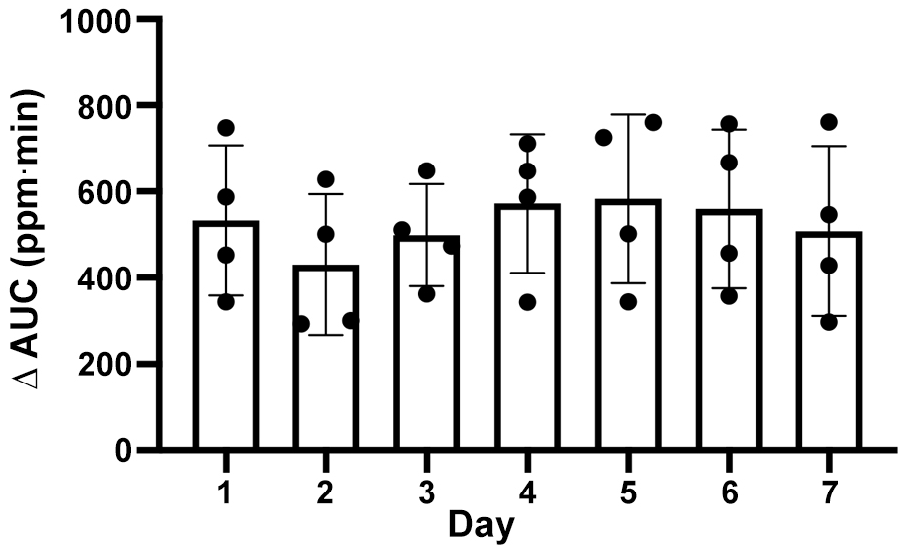
Quantification of O_2_ from lignin composites with NPs of SLS-PLGA/CaO_2_. The quantity of oxygen released from CPO lignin composites was measured using a planar O_2_ sensor attached to the bottom of a well in a 96-well plate. Area under curve (AUC) was calculated and the differential between CPO and CPOc lignin composites over 1440 min is reported. One-way ANOVA with Tukey’s *post hoc* test, no significant difference to each other; mean±SD, n=4.

### 3.3 Swelling ratios and rate of degradation were not significantly different with respect to the presence or the concentrations of NPs of SLS-PLGA/CaO_2_

Lignin composites in tissue are subject to swelling and enzymatic degradation for remodeling. Thus, we assessed the extent of swelling of lignin composites by varying the concentration of incorporated NPs of SLS-PLGA with or without CaO_2_. As shown in **Figure 3a**, the swelling ratios between CPO and CPOc lignin composites were not significantly different at both 4 and 40 mg/mL. With 40 mg/mL of NPs of SLS-PLGA in lignin composites, the swelling ratios were slightly reduced by around 7% (CPO) and 8% (CPOc) without any statistical difference. Apparently, the 10 times higher mass fraction of NPs (regardless the presence of CaO_2_) in lignin composites contributed to the swelling behavior with marginal difference. To determine the fraction of remaining lignin composites, TLS, CPO (4 and 40 mg/mL) and CPOc (4 and 40 mg/mL) lignin composites were submerged either in collagenase/CaCl_2_/serum-free medium or CaCl_2_/serum-free medium. After 24 h, less than 28% of TLS lignin composite (**Figure 3b**), 38% of CPOc lignin composite with NPs of SLS-PLGA (w/o CaO_2_) at 40 mg/mL and 23% of CPOc lignin composite with NPs of SLS-PLGA (w/o CaO_2_) at 4 mg/mL remained (**Figure 3d**) while CPO lignin composites retained around 60% at both 4 and 40 mg/mL (**Figure 3c**). All control groups (SF medium in **Figures 3b-d**) showed minimal (up to 7%) to no changes in the remaining fraction of lignin composites. The concentrations of matrix metalloproteinase (MMP) in inflamed tissue vary from patients to patients. For example, the concentration of MMP-1/18/13 in chronically inflamed dental pulp tissue is 3 ng/mL[47] and MMP-3 in rheumatoid arthritis patients in 350 ng/mL while the health controls have 75 ng/mL[48]. The concentration of collagenase type II was 0.5 U/mL (equivalent to 430 μg/mL), which is several orders of magnitude higher than the physiological conditions. Native or wounded skin has a plethora of MMPs, their inhibitors and serum proteins, leading to a tighter control of degradation and thus, we expect slower degradation of lignin composites *in vivo*. Nevertheless, we expect a certain extent of degradation of lignin composites (primarily, GelMA) *in vivo* to transiently protect NPs of SLS-PLGA/CaO_2_ from rapid, enzymatic and mechanical degradation while promoting antioxidant activity from lignosulfonate (both NPs of TLS and SLS in SLS-PLGA).

**Figure 3.**
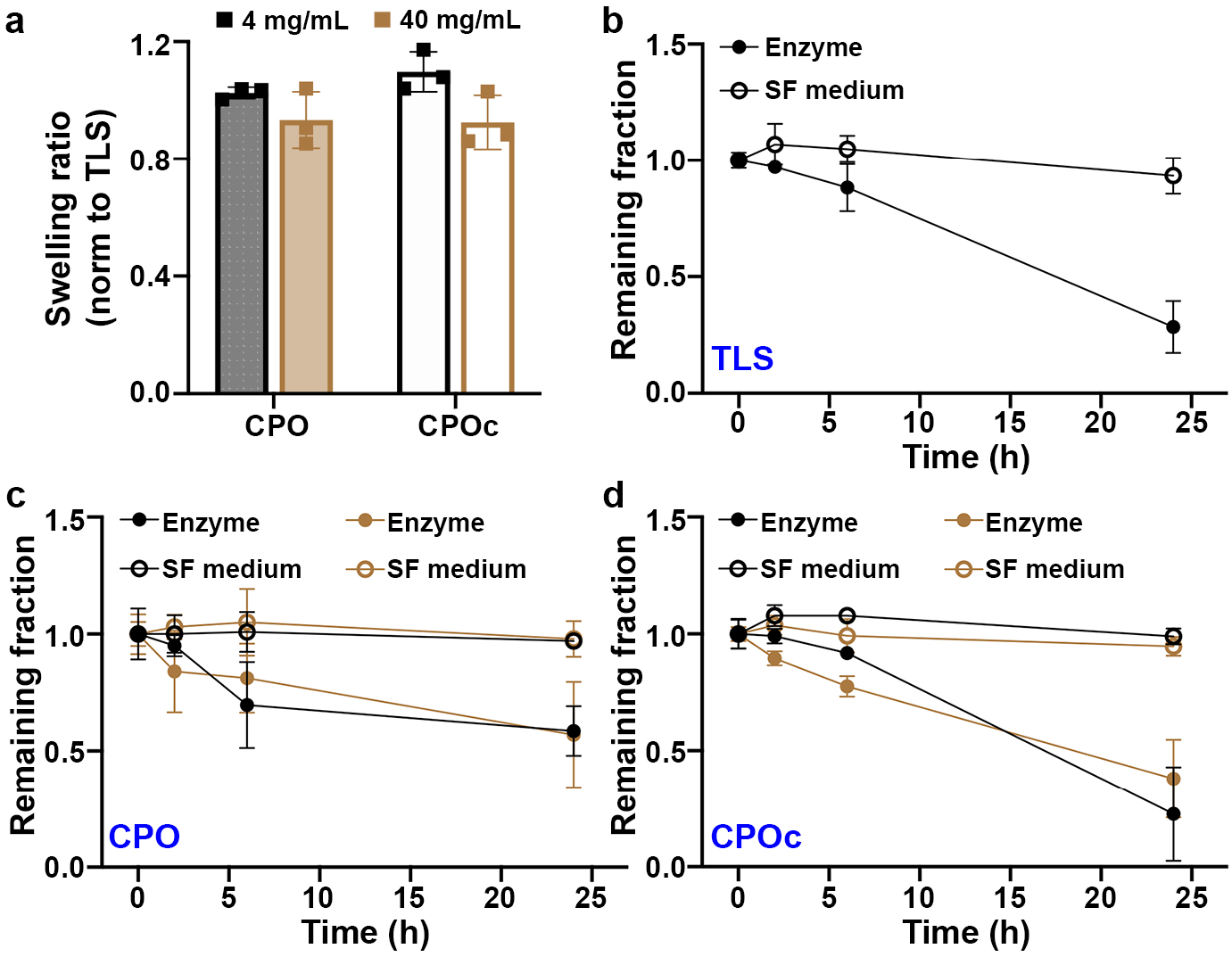
Swelling ratios and degradation of CPO and CPOc lignin composites. (a) Swelling ratios normalized to TLS lignin composites. No statistical difference between CPO and CPOc lignin composites at the same concentration and between two different concentrations at each composite type. Student’s *t*-test showed no statistically significant difference; mean±SD, n=3. Degradation of lignin composites, TLS (b), CPO (c) and CPOc (d), in the solution of 0.5 U/mL collagenase (enzyme) or serum-free medium (SF medium). In (b-d), black and brown symbols represent the concentrations of NPs (SLS-PLGA with or without CaO_2_) at 4 and 40 mg/mL, respectively; mean±SD, n=3.

### 3.4 NPs of SLS-PLGA did not alter the viscosity of the lignin composite precursors while quantity and type of NP modulated the viscoelasticity of lignin composites

Since lignin composites are subject to needle injection to the wounded areas, we assessed the mechanical properties of lignin composites before and after thiol-ene crosslinking. Viscosity of all five different types of lignin composite precursors was tested and we found no significant difference (**Figure 4a**). Apparently, crosslinked lignin composites exhibited similar viscoelasticity. However, the compressive modulus of elasticity was significantly different upon incorporating NPs of SLS-PLGA/CaO_2_ and SLS-PLGA (w/o CaO_2_). As shown in **Figure 4b** and **Table 3**, TLS lignin composites exhibited the highest modulus of elasticity. Upon incorporating NPs of SLS-PLGA/CaO_2_ at 4 or 40 mg/mL, moduli of elasticity were decreased to around 18-19 kPa. While TLS is thiolated SLS to apply thio-ene crosslinking[33], SLS in NPs of SLS-PLGA was not functionalized for crosslinking. Instead, NPs of SLS-PLGA/CaO_2_ harness CaO_2_ proximal to PLGA chains[49] while NPs of SLS-PLGA (w/o CaO_2_) were formed without any CaO_2_, which may form cavities during lyophilization. This possibly resulted in lowering the elasticity from the compression test during oscillating rheometry. Consequently, higher quantity (40 mg/mL) of NPs of SLS-PLGA (w/o CaO_2_) showed much lowered slope down to 7.27 kPa in comparison to that of CPO or TLS lignin composites. In **Figure 4c**, storage moduli of TLS and CPO lignin composites were similar to each other while those of CPOc lignin composite were significantly different from those of TLS or CPO lignin composites. However, loss tangent (G″/G′) of all lignin composites ranged from 0.01 to 0.07 (equivalent to δ (phase lag) ranging from 0.57° to 4°), indicative of well-crosslinked viscoelastic composites (**Figure 4d**). Collectively, the reduction of stiffness by non-crosslinkable NPs of SLS-PLGA/CaO_2_ is potentially significant around 40 mg/mL, thus we continued our investigation of oxygen generating capability from lignin composites with NPs of SLS-PLGA (with or without CaO_2_) at 4 mg/mL and in mouse models of wound healing.

**Figure 4.**
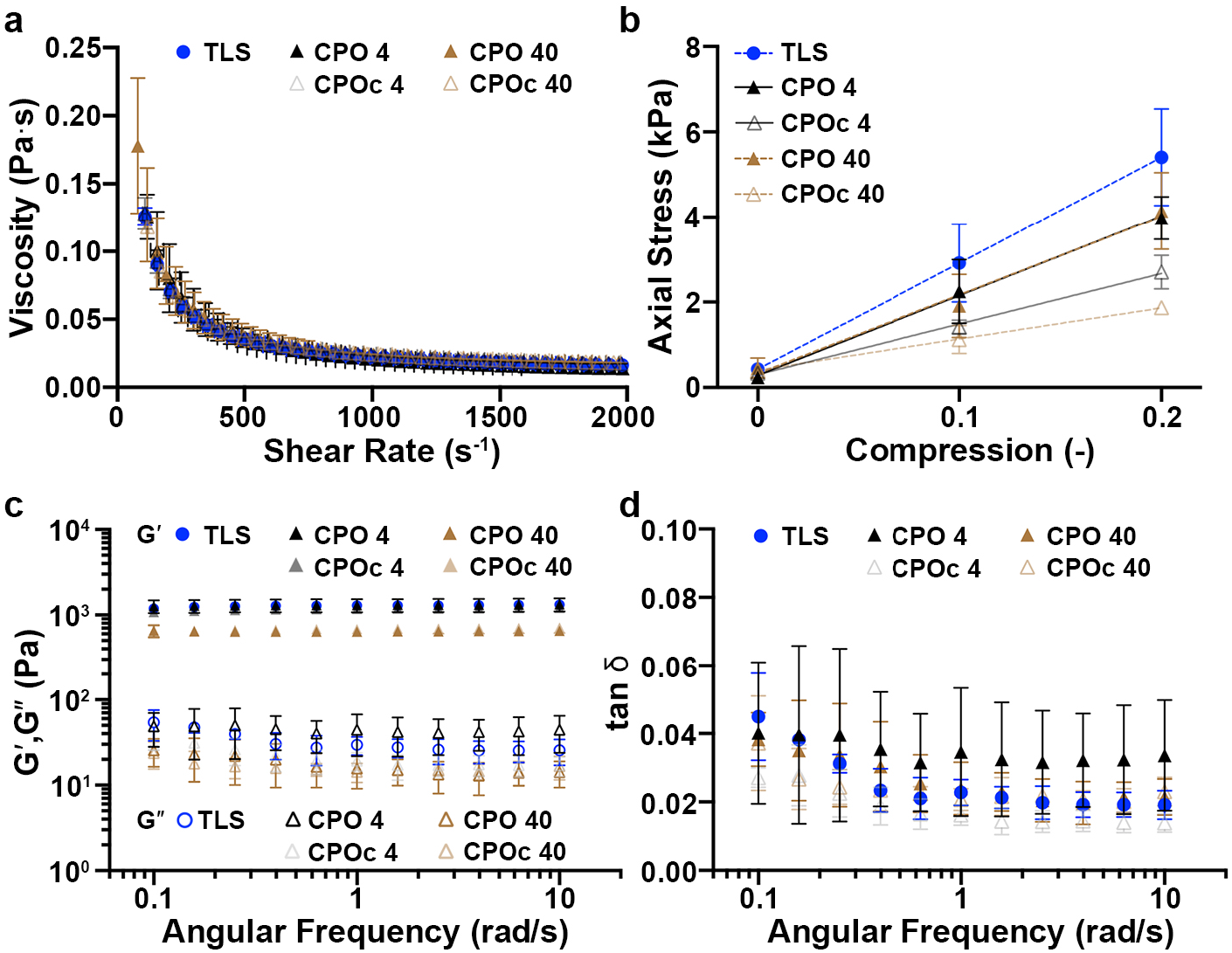
Oscillating rheometry of lignin composites. (a) Viscosity of each precursor before photo-crosslinking. (b) Axial stresses are plotted against compression varying from 0 to 20%. (c) Frequency sweeping of lignin composites. Solid and open symbols represent G′ (storage modulus) and G″ (loss modulus), respectively. (d) Loss tangent (δ) of lignin composites from 0.1 to 10 rad/s. mean±SD, n=3 for all samples.

**Table 3.** Elastic modulus estimated by the slope of axial stress vs compression.

| Lignin Composite | Slope (Elastic modulus, kPa) | Intercept (kPa) | R<sup>2</sup> |
| --- | --- | --- | --- |
| TLS | 24.9 | 0.430 | 0.8960 |
| CPO 4 mg/mL | 18.7 | 0.286 | 0.9265 |
| CPO 40 mg/mL | 18.4 | 0.329 | 0.8661 |
| CPOc 4 mg/mL | 11.8 | 0.304 | 0.9557 |
| CPOc 40 mg/mL | 7.27 | 0.409 | 0.9290 |

### 3.5 The area of granulation tissue was significantly increased with CPO lignin composites

To determine the effect of lignin composites on the progression of wound healing, wounds were harvested at day 7 post-wounding to examine re-epithelialization and granulation tissue formation (**Figure S2**). After 7 days of wounding, epithelial gap and the area of tissue granulation were assessed. Untreated wounds (UNTX in **Figure 5a**) showed no sign of re-epithelialization as expected. TLS or CPOc lignin composites in **Figure 5a** still showed the separation of GelMA matrix and tissue (insets in **Figure 5a**), while the CPO lignin composite enhanced integration of the lignin composite into the wounded tissue over 7 days. There was no difference of the rate of the wound closure, as determined by epithelial gap (the distance between the arrows), between UNTX and TLS (5.0±1.4 vs. 5.8±1.2 mm, p>0.05), CPOc (5.0±1.4 vs. 4.7±0.8 mm, p>0.05), or CPO (5.0±1.4 vs. 4.4±0.7 mm, p>0.05) lignin composite treated mice (**Figure 5b**). However, we observed significantly increased granulation tissue, between UNTX and TLS (1.4±0.2 vs. 2.1±0.3 mm, p=0.01) or CPOc (1.4±0.2 vs. 2.2±0.3 mm, p=0.01) lignin composite treated mice, especially with CPO lignin composite treatment (1.4±0.2 vs. 3.0±0.5 mm^2^, p<0.01) (**Figure 5c**), which is indicative of healthy healing progress.

**Figure 5.**
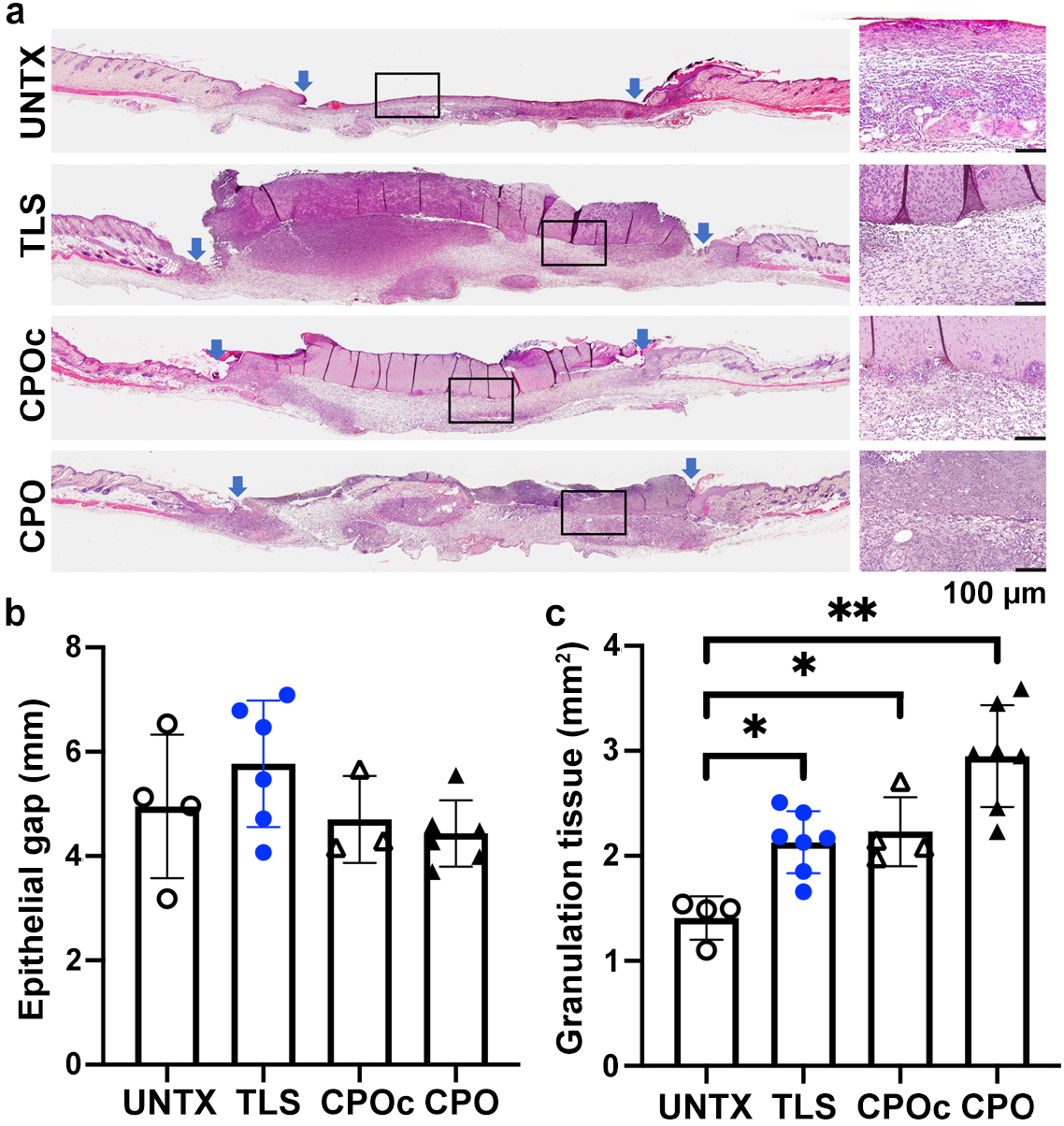
Morphometric analysis of wounds treated with lignin composites at 7 days after surgery. Wounds in WT C57BL/6N mice were treated with lignin composites (a) to measure epithelial gap (b) and granulation tissue area (c). In (a), boxed areas are provided in inset images at higher magnification of hematoxylin (blue, nuclei) and eosin (red, ECM and cytoplasm) staining. Scale bar, 100 μm. One-way ANOVA with Kruskal-Wallis test followed by Dunn’s multiple comparison test. *p<0.05 and **p<0.01. 4≤n≤7, mean±SD. Details of compositions of UNTX, TLS, CPOc and CPO are in **Table 1**.

### 3.6 Increased vascularity was promoted by the CPO lignin composite over 7 days

In addition to the increase of granulation tissue formation, neovascularization is promoted by the CPO lignin composites. Quantification of neovascularization (**Figure 6a**) showed that UNTX has a significantly higher number of CD31^+^ ECs while the formation of blood vessel was delayed at day 7 (**Figure 6b**). Staining with the EC marker (CD31) revealed a greater number of individual CD31^+^ cells that are not associated with vessels in UNTX wounds (**Figure 6b**) per HPF than TLS-(36.4±19.8 vs. 11.6±6.3%, p<0.05), CPOc-(36.4±19.8 vs. 4.2±4.3%, p<0.01) or CPO-(36.4±19.8 vs. 15.1±7.7%, p<0.05) lignin composite-treated wounds. However, there were significantly more vessel lumens (**Figure 6c**) per HPF in the CPO lignin composite-(17.3.2±7.6 vs. 17.3±7.5 vessels/HPF, p<0.05) treated wounds as compared to UNTX wounds.

**Figure 6.**
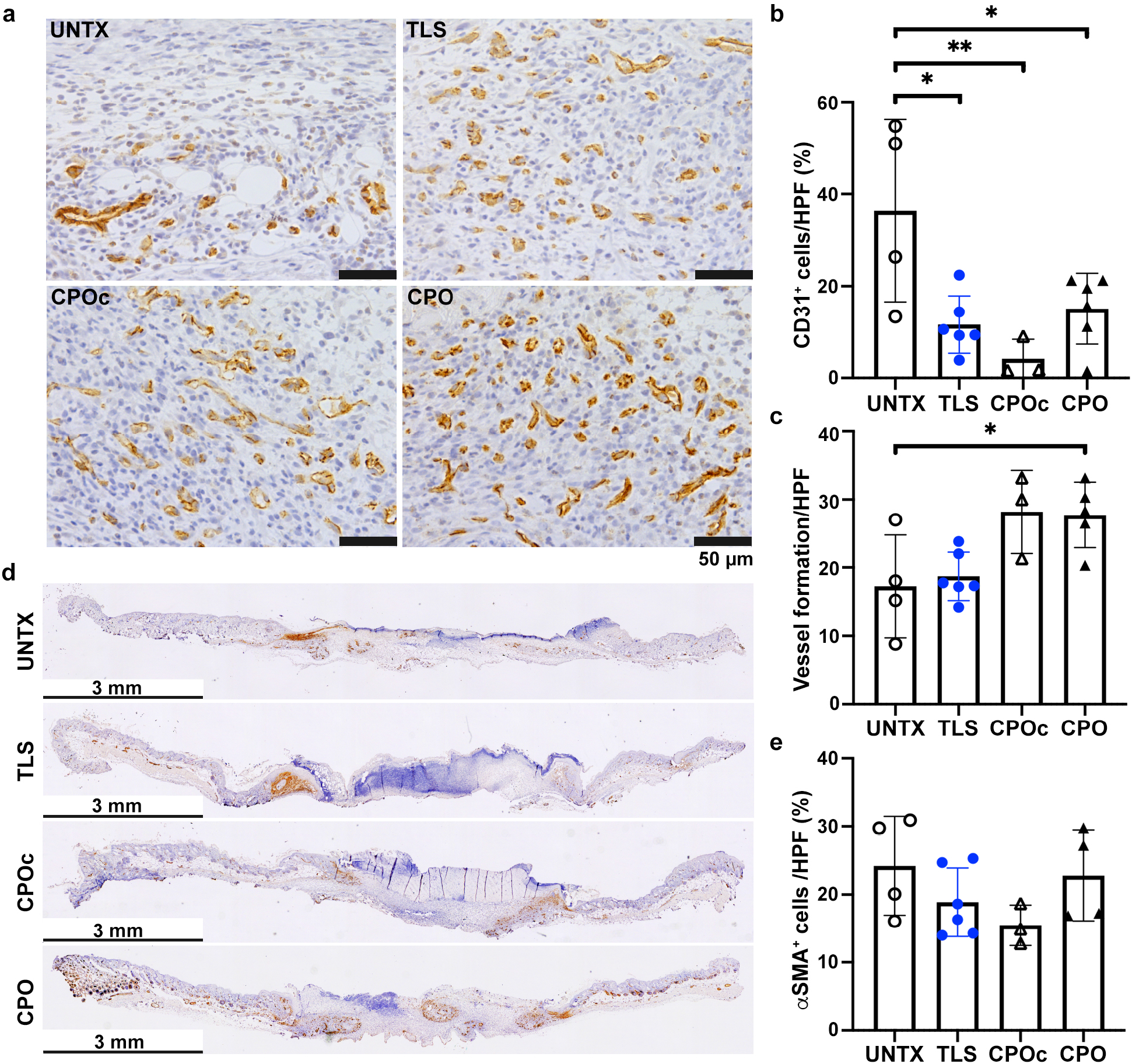
Assessment of vascularization in the wounds treated with lignin composites at 7 days after surgery. Wounds in WT C57BL/6N mice were treated with lignin composites to assess the extent of vascularization with immunostaining (CD31, brown) and hematoxylin counterstaining (blue, nuclei). CD31^+^ cells (b) and vessel formation (c) per HPF were quantified. The infiltration of αSMA^+^ cells to wounds was visualized (d) and quantified per HPF (e). Scale bars, 50 μm in (a) and 3 mm in (d), respectively. One-way ANOVA with Tukey’s *post hoc* tests, **p<0.01 and *p<0.05. 3≤n≤6, mean±SD. Details of compositions of UNTX, TLS, CPOc and CPO are in **Table 1**.

As shown in **Figure 6d**, αSMA^+^ cells (brown staining) were limited to the edge of wounds in UNTX and TLS lignin composites. CPO or CPOc lignin composites showed more αSMA^+^ cells in the wounds. However, the CPO lignin composite showed a higher numbers of blood vessel formation both at edge and middle of wounds (**Figure 6d**). Review of the literature demonstrates that the expression of αSMA implicates the development of vasculature[50, 51] and is present in pericytes on capillaries[52]. While the extent of difference at either edge or middle sections was not significantly different (**Figure S3**), the overall trend was an increase with the CPO lignin composites (**Figure 7e**). We observed similar trends in the number of CD31^+^ cells (**Figure 7b**) and αSMA^+^ cells (**Figure 7e**). The expression of αSMA^+^ by myofibroblasts (myoFB) is also shown to underlie tissue regeneration in the skin, and the number of αSMA^+^ cells decrease as the regeneration process is completed[53]. Thus, neovascularization is promoted, and the wound healing is still in progress by day 7 with the injection of the CPO lignin composite.

**Figure 7.**
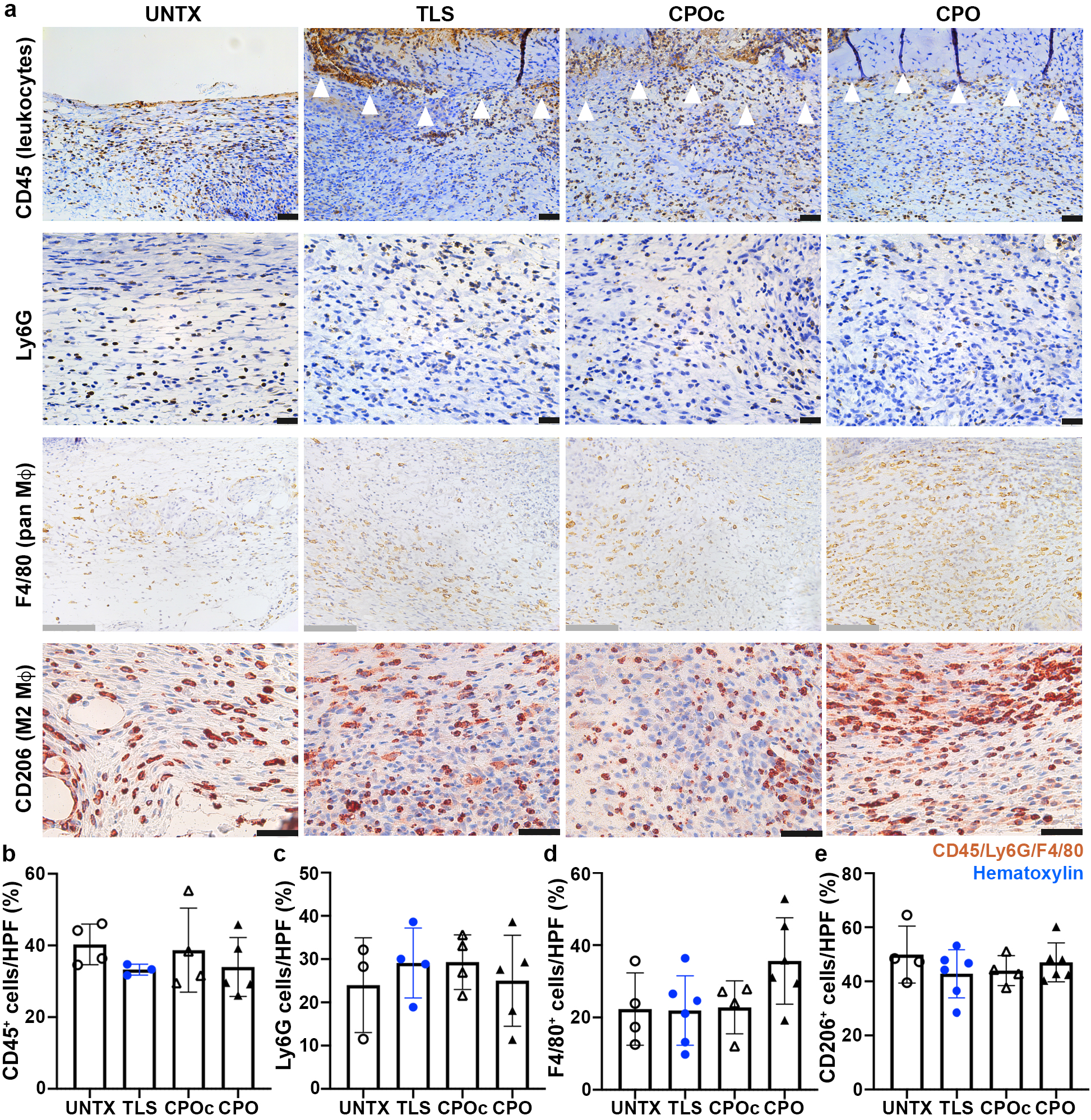
Assessment of inflammatory responses from the wounds treated with lignin composites at 7 days after surgery. Wounds in WT C57BL/6N mice were treated with lignin composites to assess the extent of inflammatory responses (a) with immunohistochemical staining and hematoxylin counterstaining (blue, nuclei). CD45^+^ leukocytes (b), Ly6G^+^ monocytes, granulocytes, and neutrophils (c), F4/80^+^ pan Mφ (d) and CD206 M2 Mφ (e) per HPF were quantified. In (a), black scale bars, 50 μm and grey scale bars, 125 μm, respectively. 3≤n≤6, mean±SD. Details of compositions of UNTX, TLS, CPOc and CPO are in **Table 1**.

### 3.7 Lignin composites did not cause significant inflammatory responses over 7 days

Following the quantification of granulation tissue formation and neovascularization, inflammatory responses were assessed by immunohistochemical staining (**Figure 7**). CD45 is expressed by common leukocytes except platelet and red blood cells. Ly6G is expressed by monocytes, granulocytes and neutrophils. F4/80 is used to identify murine tissue macrophages. CD206 is normally expressed on the alternatively activated, anti-inflammatory (M2) macrophages (Mφ). As shown in **Figure 7a**, the interface between wounds and lignin composites does not show (indicated by white wedges) significant infiltration of CD45^+^ cells. Recently, the stiffness of GelMA is known to directs macrophage phenotypes. Soft GelMA matrix (stiffness less than 2 kPa) is more favorable for priming macrophages toward M2 phenotypes with a decreased capacity for spreading in comparison to stiff (over stiffness 10 kPa) GelMA matrix[54]. The lignin composites are in the range of 1-2 kPa of storage modulus (**Figure 4c**), thus likely priming macrophage toward anti-inflammatory (M2) phenotypes. Over 7 days, no significant changes in the expression of the four markers were noticed. There was no significant difference in percentage of CD45^+^ (pan-leukocyte) cells/HPF across all tested lignin composites when compared to UNTX wounds (UNTX 40.3±5.7 vs. TLS 33.3±1.5 vs. CPOc 38.7±11.7 vs. CPO 34.0±8.2%, p>0.05) (**Figure 7b**). A similar trend was observed when looking at monocyte, granulocyte, and neutrophil infiltration as determined through Ly6G staining (**Figure 7c**). There were no significant differences in the percentage Ly6G^+^ cells/HPF among treatment groups (UNTX 24.0±11.0 vs. TLS 29.1±8.1 vs. CPOc 29.3±6.3 vs. CPO 25.1±10.5, p>0.05). In particular, alternatively-activated M2 macrophages are important mediators of successful wound healing[55]. F4/80 and CD206 staining (**Figure 7c** and **d**) showed that UNTX wounds had no statistically significant differences in the levels of F4/80^+^ cells per HPF (UNTX 22.3±10.0 vs. TLS 21.9±9.6 vs. CPOc 22.8±7.3 vs. CPO 35.6±12.0%, p>0.05) or CD206^+^ cells per HPF (UNTX 49.9±10.5 vs. TLS 42.8±8.9 vs. CPOc 44.0±5.5 vs. CPO 47.0±7.2%, p>0.05).

### 3.8 While collagen content was not significantly modulated, the wounds treated with CPO lignin composite exhibited mature collagen architecture over 30 days

Tissue sections were prepared to assess collagen architecture and content over 30 days of healing process from trichrome staining. In rodents, normal tissue has a reticular collagen pattern, whereas the collagen in scar tissue forms large parallel bundles approximately perpendicular to the basement membrane[56]. The CPO lignin composite exhibited more pronounced, bundled mesh network as opposed to relatively straight fibers found in TLS or CPOc lignin composites (**Figure 8a**). Despite qualitative difference of collagen architecture, the quantity of collagen content over 30 days becomes similar between treatment groups (**Figure 8b**). No differences were observed in the overall collagen density between UNTX and TLS-(187.0±23.8 vs. 170.0±26.4 pixels/HPF, p>0.05), CPOc-(187.0±23.8 vs. 179.6±21.5 pixels/HPF, p>0.05), or CPO-(187.0±23.8 vs. 180.9±29.4 pixels/HPF, p>0.05) lignin composite-treated wounds. However, reduction of collagen content in a scar has less impact on scarring outcome compared to the structure of matrix deposited in the dermis[57]. Further, wounds treated with the CPO lignin composite left minimal scar as evidenced in photographs taken at 30 days after surgery (**Figure S4**).

**Figure 8.**
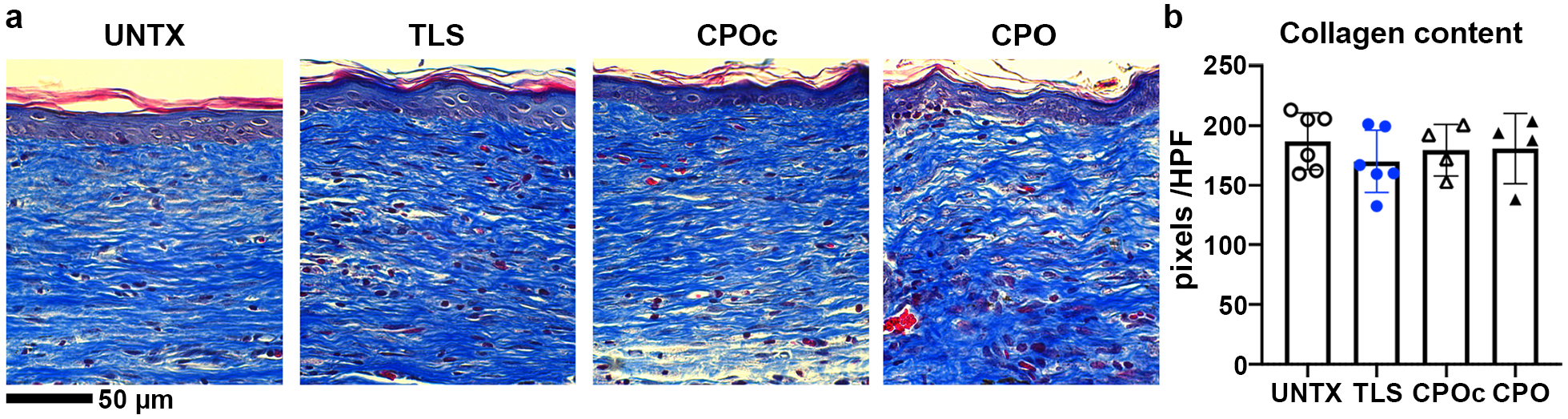
Scar assessment in the wounds treated with lignin composites at 30 days after surgery. Wounds in WT C57BL/6N mice were treated with lignin composites to assess the collagen content with trichrome (blue, collagen) (a). Collagen content was quantified per HPF using color moment thresholding in ImageJ (b). Scale bar, 50 μm. 3≤n≤6, mean±SD. Details of compositions of UNTX, TLS, CPOc and CPO are in **Table 1**.

## 4. DISCUSSION

### 4.1 Application of polyphenolic lignin polymer to wound healing and tissue engineering applications is increasing

Application of lignin is ever increasing in a number of industrial and biomedical applications, including UV absorption[58], Cu^2+^ chelating in water[59], skin protectants[60], wound dressing with chitosan[61], self-healing smart biomaterials[62], durable and repeatable adhesive[63], antibacterial agent[64, 65] with metallic NPs[66], anti-proliferative effect to cancer cells without cytotoxicity[67]. In tissue engineering, lignin composites can be applied to enhance mechanical properties with good protein adsorption capacity and wound compatibility[68], to provide 3D microenvironments of mesenchymal stromal cell culture[61], to confer anti-inflammatory properties by reducing gene expression of inducible nitric oxide synthase (iNOS) and IL-1β of inflamed mouse Mφ[69], to apply for wound-dressing with biocompatibility and nontoxicity[70], to improve mechanical properties and wound healing capabilities on incision-induced Sprague-Dawley rats[71], to provide nanofibers for the cultures of human dermal fibroblasts (hdFBs)[72], to promote localized and prolonged antioxidant capabilities for wound healing applications[73] and to achieve enhanced mechanical properties and viability of hdFBs for direct ink writing 3D bioprinting[74].

Nevertheless, toxicological studies of lignin have only carried out through simple cytotoxicity testing, which cannot accurately simulate the human body environment. Thus, the biological effects of lignin with animal models should be tested to assess the long-term stability and potential negative effects of lignin and lignin-derived biomaterials. In addition to toxicological studies, another difficulty and key point for the development of lignin-based biomaterials is the heterogeneity of lignin. Although lignin displays good potential for biomedical applications[75], its broad distribution of molecular weight and extremely complex structures represent big hurdles for deep studies, standardization, and scale-up. Thus, lignin chemistry and de/polymerization techniques still need to be continuously developed[76–79].

### 4.2 CaO_2_ from the CPO lignin composite is applicable for oxygen generating compounds for tissue regeneration

The injectable lignin composite can be a unique biomaterial platform to confer antioxidation by TLS and oxygen generation from NPs of SLS-PLGA/CaO_2_ without incorporating additional enzymes. Incorporation of CaO_2_ into NPs of SLS hydrophilic outer layer and PLGA hydrophobic core[35] allows CaO_2_ to be complexed with PLGA. CaO_2_ is a white to yellowish powder which has been extensively applied in water treatment, seed disinfection and food processing[80]. When reacted with water at pH lower than 12 (**Figure S5A**), CaO_2_ can be decomposed into hydrogen peroxide, hydroxide ions and carbonate, and the generated hydrogen peroxide further decomposes into highly reactive superoxide and hydroxyl radical[81]. Catalase can decompose the intermediate hydrogen peroxide into water and oxygen. While catalase is an enzyme found in the blood and liver of mammals, this enzyme has to be available *in situ*, not from peroxisome to prevent damages from ROS. This intermediate step eliminates any potential cytotoxic ROS[82]. Without catalase, the cytotoxic byproduct H_2_O_2_ may lead to cell damage.[83] Although the actual role of catalase in the oxygen-release process is unclear, decomposition is suggested to take place through the Modified Fenton chemistry – dissociation of H_2_O_2_ to OH radicals with Fe^2+^/^3+^ [84]. These oxidants react with everything within diffusion limit layer while their half-lives are short. In wound healing, the availability of catalase is limited. Even though the level of expression of catalase mRNA is not changed during wound healing[85], the protein level of catalase is decreased[86]. For example, catalase concentration is reported to be in the range of 50 – 100 U/mL in the oxygen generating gelatin section[87] or 1 mg/mL in the matrix of polycaprolactone (PCL)[88]. In the mixture of CaO_2_, catalase and either SLS or TLS, the dissociation of H_2_O_2_ is predominantly facilitated by catalase and partly by TLS and to less extent by SLS (**Figure S5B**). The native antioxidant properties of SLS or TLS NPs (**Figure S6A**) is maintained at levels of 80% or above of the native antioxidant (*L*-ascorbic acid) in the presence of up to 50 μg/mL of CaO_2_ (**Figure S6B**).

Several cases of CaO_2_ incorporation to biomaterials are reported to date. Injectable cryogels are formed with methacrylated hyaluronic acid and CaO_2_[89]. Cumulative release of H_2_O_2_ is up to 468 μmol over 3 h. Microparticles (MPs) are formed with the encapsulation of CaO_2_ in PCL, then formed composites with GelMA[90] and release O_2_ up to 5 weeks[91]. Over 2 weeks, 2.5 μM of H_2_O_2_ is released without the formation of MPs while 1.0 μM with the formation of MPs. With 5 to 20 mg/mL of CaO_2_, the cumulative release of O_2_ is 25% without MP and up to 30% with MP. Some cases include catalase (100 U/mL) and CaO_2_ (up to 30 mg/mL)[92, 93]. GelMA (50 mg/mL)-CaO_2_ (30 mg/mL) with catalase (100 U/mL) releases O_2_ up to 5 days[92]. Another type of hydrogel utilizes Ca^2+^ from CaO_2_ (up to 10 mg/mL) to crosslink gellan gum (anionic polysaccharide) with catalase[94]. In the matrix of silk/keratin/gelatin, CaO_2_ up to 200 mg/mL release O_2_ over 2 weeks with additional antibacterial properties for urethral tissue engineering applications[95]. Thiolated gelatin (27 to 63 mg/mL) with CaO_2_ (25 to 100 mg/mL) and catalase (2000-5000 U/mg) forms hyperbaric oxygen generating biomaterials.[96] The question is how much CaO_2_ is required to efficiently deliver O_2_ to tissue microenvironments without causing cytotoxicity. CaO_2_ concentrations higher than 10 mg/mL exhibit cytotoxicity to 3T3 FBs^78^. In addition, due to low solubiliuty in water, it is difficult to disperse CaO_2_ in aqueous buffers. Further, Ca^2+^ released from these composites influence cell cytotoxicity and bacterial adhesion[97].

To assess the kinetics of O_2_, *in situ* oxygen measurement with patch sensors are used[88, 96, 98]. In the case of measuring concentrations of H_2_O_2_, Cu(II)-neocuproine spectrophotometric method[99] or free radical analyzer[100] are used. To delineate the difference, the release kinetics of O_2_ and H_2_O_2_ from CaO_2_ in water with varying pH and temperature is probed and modeled[101]. The release of H_2_O_2_ follows pseudo-zero order (PZO) reaction. Once CaO_2_ is dissolved in water, Ca^2+^, O_2_^2-^ and H^+^ are formed. Of these, 2H^+^ and O_2_^2-^form H_2_O_2_. With increase of H^+^ (decrease of pH), the solubility of CaO_2_ is increased. Temperature has minimal to no effect in the kinetics. In contrast, the release of O_2_ follows pseudo-first order (PFO) reaction, meaning that the concentration of CaO_2_ directly affects the dissociation of CaO_2_ to O_2_. With the decrease of pH, the release of O_2_ is decreased. With the increase of temperature, the release of O_2_ increases. From this study of kinetics, H_2_O_2_ is not necessarily a precursor of O_2_.

### 4.3 Locoregional delivery of NPs for antioxidation and oxygen generation could be beneficial to promote wound healing

Our previous work[33] showed that the antioxidation capacity of TLS attenuated the expression of *COL1A1*, *ACTA2*, *TGFB1*, and *HIF1A* and resulted in the similar extent of expression of the fibrotic markers comparable to proliferative LS hdFB phenotypes. To further confirm that the attenuation of fibrotic phenotypes of hdFBs by TLS lignin composite, we analyzed 84 genes associated with angiogenesis, ECM production and oxidative stress using a PCR array. We hypothesized that exposure of LS and HS hdFBs to TLS lignin composite attenuates these angiogenesis, ECM production and oxidative stress responses of HS to the levels comparable to those of LS. As shown in **Figure S7**, the expression of fibrotic genes in HS hdFBs on lignin composites was significantly altered to mimic those in LS. These results and the previous works[33] suggest that addition of lignosulfonate to the microenvironment of wound healing may attenuate fibrotic responses during tissue repair. Further, we are also inspired to include sustained supply of oxygen *in vivo* for enhanced angiogenesis (**Figure 6**)[102] and ECM production (**Figure 8**) using a biomaterial platform based on lignosulfonate.

Although WT mice are programmed to heal physiologically with minimal alterations in perfusion and in the presence of ROS, we did see improvement in our measures of wound healing (**Figure 5**) and vessel formation (**Figure 6**) without significant inflammation (**Figure 7**). While the quantity of released oxygen (around 700 ppm per day) is suboptimal yet, the formulation of lignin composites investigated here is rather an effective suboptimal dose of oxygen without causing excessive scarring by VEGF (vascular endothelial growth factor) stimulation[103]. Thus, the enhanced wound healing by CPO lignin composite is the product of enhanced vascularization *and* granulation tissue formation.

The translatable benefit of the lignin composite may be more readily apparent in disease states such as diabetic wound healing and/or with infection. The advanced glycation end products (AGEs) present in diabetic tissue generate ROS leading to FB apoptosis via the NLRP3 (NOD-, LRR- and pyrin domain-containing protein 3) signaling pathway, thus impairing wound repair[104]. Some systemic oral anti-hyperglycemics currently used to treat diabetes mellitus also have antioxidant and beneficial wound healing effects. Metformin, for example, is a commonly used medication used for glucose control in diabetes mellitus patients which has antioxidant effects and demonstrated benefits on angiogenesis and wound closure in diabetic mice[105]. However, as previously mentioned, pure systemic antioxidants have a poor enteric absorption profile. Locally applied antioxidant hydrogels have been demonstrated to accelerate diabetic wound healing while promoting M2 macrophage differentiation and reducing IL-1β production[106]. Further, topical oxygen delivery to ischemic wounds is shown to accelerate healing and promote granulation tissue formation[107]. The strategy of combining these dual acting, wound-beneficial attributes into one local therapy for pathologic wounds has the potential to greatly benefit the wound healing in at-risk patients.

## 5. CONCLUSIONS

Lignin has been applied to many different tissue engineering applications. Wound healing applications of lignin has been somewhat limited to doping lignin NPs into a matrix of biomaterials. First, we functionalized lignosulfonate to form dual acting lignin composites with TLS and SLS-PLGA NPs. We designed NPs of SLS-PLGA including CaO_2_ to deliver O_2_ without adverse effect of tissue oxygenation. The quantity of released O_2_ from the CPO lignin composite was around 700 ppm/day at least for 7 days. Mechanical properties of lignin composites were not significantly different, but the stiffness set for the *in vivo* applications was amenable to direct M2-like macrophage phenotypes. Swelling and enzymatic degradation of lignin composites *in vitro* were verified before the animal experiments. The area of granulation tissue formation was significantly increased by the CPO lignin composite while the rate of wound closure was not significantly different by 7 days after surgery. However, the CPO lignin composite increased neovascularization as evidenced by significantly increased blood vessel formation and by the infiltration of αSMA^+^ cells to the wound treated with the CPO lignin composite over 7 days. In scar assessment, while we did observe no difference in the overall collagen density, the collagen architecture treated by the CPO lignin composite exhibited more pronounced, bundled mesh network and left minimal scar. Thus, soft matrix with antioxidation (conferred by TLS) and synergistic oxygen release (conferred by SLS-PLGA/CaO_2_ NPs) could be applied to wound healing applications with enhanced tissue granulation, vascularization and maturation of collagen architecture without significant inflammatory responses. Further optimization of the dual component lignin composite is in progress, aiming to significantly accelerate diabetic wound healing with infection.

## ABBREVIATIONS

ANOVA: analysis of variance
AUC: area under curve
DLS: dynamic light scattering
ECs: endothelial cells
ECM: extracellular matrix
FBs: fibroblasts
FFPE: formalin fixed paraffin embedded
HPF: high powered field
hdFBs: human dermal fibroblasts
iNOS: inducible nitric oxide synthase
LAP: lithium phenyl-2,4,6-trimethylbenzoylphosphinate
GelMA: methacrylated gelatin
MPs: microparticles
myoFB: myofibroblasts
NPs: nanoparticles
PIXE: particle-induced x-ray emission
PLGA: poly(lactic-co-glycolic) acid
PDMS: polydimethylsiloxane
PFO: pseudo-first order
PZO: pseudo-zero order
ROS: reactive oxygen species
SLS: sodium lignosulfonate
TLS: thiolated lignosulfonate
TCEP HCl: tris(2-carboxyethyl)phosphine hydrochloride
WT: wildtype

## CRediT AUTHOR STATEMENT

Swathi Balaji: Conceptualization, Methodology, Validation, Formal analysis, Respurces, Data curation, Writing – original draft, Writing - review & editing, Visualization, Supervision, Project administration, Funding acquisition

Walker D. Short: Validation, Formal analysis, Investigation, Data curation, Writing – original draft Benjamin W. Padon: Formal analysis, Investigation, Data curation

Jorge A. Belgodere: Investigation, Data curation

Sarah E. Jimenez: Investigation, Data curation

Naresh T. Deoli: Methodology, Software, Investigation, Data curation

Anna C. Guidry: Investigation, Data curation

Justin C. Green: Investigation, Data curation

Tanuj J. Prajapati: Investigation, Data curation

Fayiz Farouk: Investigation, Data curation

Aditya Kaul: Investigation, Data curation

Dongwan Son: Investigation, Data curation

Olivia S. Jung: Investigation, Data curation

Carlos E. Astete: Resources, Investigation, Data curation

Myungwoong Kim: Methodology, Resources, Funding acquisition

Jangwook P. Jung: Conceptualization, Methodology, Validation, Formal analysis, Resources, Data curation, Writing – original draft, Writing - review & editing, Visualization, Supervision, Project administration, Funding acquisition

## ACKNOWLEDGMENTS

The authors acknowledge the support from Gill Plastic Surgery of Houston (SB), Wound Healing Society Foundation FLASH and 3M Award (SB), Clayton Award from Department of Surgery, Texas Children’s Hospital (SB), John S. Dunn Foundation Collaborative Research Award (SB), the National Institutes of Health T34GM136452 (SEJ), LSU Discover Undergraduate Research Program (ACG), the National Research Foundation of Korea NRF-2021R1A2C1093999 (DS and MK), the National Science Foundation EPSCoR Track 2 RII, OIA 1632854 (JAB and JPJ) and the National Science Foundation CAREER DMR 2047018 (JPJ). The authors sincerely acknowledge the technical support received from Ashkan YekrangSafakar and Kidong Park (LSU).

## DATA AVAILABILITY

The raw data from this study are available from the authors upon request.

