## Supplementary Information for "Engineering Antioxidant and Oxygen-Releasing Lignin Composites to Promote Wound Healing"

### SUPPLEMENTARY METHODS

Alkene incorporation reaction in gelatin

Alkene was incorporated into gelatin by the coupling reaction of gelatin with methacrylate acid (MA). Gelatin (1.00 g) was dissolved in 10.0 mL of PBS (pH = 7.4) at 50°C. Separately, a solution of MA (0.10 g), *N*-ethyl-*N′*-(3-(dimethylamino)propyl)carbodiimide (EDC; 0.178 g), *N*-hydroxysuccinimide (NHS, 0.107 g) and dimethyl sulfoxide (DMSO, 2.00 mL) was prepared by stirring at 40°C for 30 min. The flask containing the gelatin solution was placed in an oil bath at 50°C, and the other solution of MA, EDC, NHS and DMSO was added dropwise. The mixture was further stirred at 50°C for 1.5 h. The solution was cooled down to room temperature, followed by dialysis in water at 40°C for a week. The resulting solution was lyophilized to yield alkene-incorporated gelatin powder sample. The coupling efficiency of MA to gelatin was assessed using $^1$H NMR comparing the phenyl peak (a) and the lysine peak (b) (**Figure S1**). Upon coupling reaction, the development of two distinct peaks between 5 to 6 ppm were observed in $^1$H NMR spectrum, referring to the two protons in alkene group of MA attached to the lysine of gelatin. Additionally, the gelatin phenylalanine (Phe) group remained untouched during the reaction and allowed us to determine the mmol/g ratio of the GelMA. From literature[1], native porcine gelatin contains 15.5 mol/$10^5$ g of Phe. Combining this information with the ratio of Phe:MA, the mmol/g concentration of MA was calculated using the following equation: MA concentration (mmol/g) = (15.5 mol/$10^5$ g) $\times$ (1000 mmol/mol) $\times$ ($I_{phe}/I_{alkene}$) (intensity ratio of phenyl to alkene). The batch presented in **Figure S1** showed the $I_{phe}/I_{alkene}$ of 1:1.905 ([intensity from five protons of Phe/5] / [intensity of two protons of alkene/2]) resulting in a MA concentration of 0.295 mmol/g. This value also can be utilized to estimate the degree of substitution with a ratio of 1:2.20 ([intensity from five protons of Phe/5] / [intensity of two protons on carbon adjacent to $NH_2$ (c) in Lys/2]) of Phe:Lys of the pristine gelatin (i.e. [reacted MA upon the substitution]/[reactive amine site (Lys) in pristine gelatin] $\times$ 100 = 1.905/2.20 $\times$ 100 = 87%), suggesting that the modifications made in our synthesis route produced GelMA with around 87% substitution.

Cultures of high and low scar normal human dermal fibroblasts

Skin tissue was collected from abdominoplasty patients who provided written informed consent as part of a protocol approved by the Institutional Review Board of Texas at Baylor College of Medicine in accordance with the Declaration of Helsinki. A biobank of scar and matched normal uninjured skin tissue was obtained from abdominoplasty patients controlled for sex, age, ethnicity, surgery type, indication, wound site, and comorbidities. These skin tissues were grouped into 'low scarring' and 'high scarring' phenotypes based on the evaluation of their existing c-section scars using Vancouver Scar Scale (VSS). Skin obtained from patients with 1-3 score on VSS were categorized as 'low scarring' phenotype (N) and patients with 6-9 score were grouped into 'high scarring' phenotype (S). For each group, n=3 skin tissues were pooled. Fibroblasts (FBs) were isolated from scar and matched normal skin cohorts and were called 'low scarring' (LS) and 'high scarring' (HS) normal fibroblasts based on their VSS phenotype and patient demographics. Only LS-normal and HS-normal human dermal FBs (hdFBs) were used for cell studies, and they are denoted HS and LS for the rest of the studies. Both LS and HS hdFBs were maintained in DMEM (Cat# 10567-014-500mL, Thermo Fisher Scientific, Waltham, MA, USA) supplemented with 10% fetal bovine serum (FBS, Cat# 35-015-CV, Corning, Corning, NY, USA), Penicillin/Streptomycin (P/S, Cat# 15140-122, Thermo Fisher Scientific, Waltham, MA, USA), and antibiotic/antimycotic (anti/anti, Cat# 15240-062, Thermo Fisher Scientific, Waltham, MA, USA). Only passages 8-10 were used for studies and cells were harvested at 80-90% confluency by applying TrypLE Express (Cat# 12604-021, Thermo Fisher Scientific, Waltham, MA, USA) and pelleted at 350 g for 5 min. Supernatant was removed and hdFBs were gently resuspended in cell culture media and counted. Media was changed every other day.

PCR array

To determine the effect of lignin composite on their expression of fibrotic phenotype of hdFBs, LS and HS hdFBs were seeded onto composites at 50,000 cells/cm$^2$ and maintained for 24 h in a $CO_2$ incubator (5% $CO_2$/37°C). For the fibrosis array, the concentrations of GelMA and LAP were fixed at 80 and 5 mg/mL, respectively. At specified time points, cell-seeded composites were removed and incubated with TrypLE Express to extract RNA by following the procedure from the PureLink RNA kit (Cat# 12183018A, Thermo Fisher Scientific, Waltham, MA, USA). Extracted RNA was evaluated for quantity and purity using a Take3 Micro-Volume plate and Cytation3 spectrophotometer (Biotek, Winooski, VT, USA). cDNA was reverse-transcribed using High-Capacity RNA-to-cDNA kit (Cat# 4387406, Thermo Fisher Scientific, Waltham, MA, USA) following the protocol from the manufacturer. We also used the RT2 Profiler PCR Array human fibrosis kit (Cat# PAHS-120-ZE-4, Qiagen, Hilden, Germany) to detect the expression of 84 fibrosis-related genes. cDNA (400 ng) made with RT2 first strand kit (Cat# 330404, Qiagen, Hilden, Germany) were used with RT2 SYBR Green qPCR Mastermix (Cat# 330501, Qiagen, Hilden, Germany) to run qRT-PCR on Bio-Rad CFX 384 real-time system. Data were analyzed by Qiagen RT2 Profiler PCR array data analysis software. GAPDH was used as a housekeeping gene, and ΔΔCT values for the genes were analyzed. Principal component analysis (PCA) and hierarchical clustering (HC) were performed and visualized with an open source machine learning software (Orange)[2].

Quantification of $H_2O_2$ release from $CaO_2$

The amounts of $H_2O_2$ released from $CaO_2$ were quantified using a Quantitative Peroxide Assay Kit (Pierce, Cat#23280) using the ferrous ion ($Fe^{2+}$) oxidation xylenol orange assay[3]. $CaO_2$ was dissolved in PBS at different pH. $CaO_2$ was also dissolved in PBS containing catalase (100 U/mL, Fisher Scientific, Cat#S25239A), SLS or TLS (3 mg/mL)[4] at 37°C. An aliquot of 100 µL of working reagent was added to the collected sample of 10 µL and incubated at room temperature for 15 min. The absorbance was measured at 595 nm using Cytation3 spectrophotometer (Biotek, Winooski, VT, USA).

Impact of the presence of $CaO_2$ on antioxidation capacity of SLS and TLS

The antioxidant activity of SLS and TLS was evaluated using the 2,2-dipehnyl-1-picrylhydrazyl (DPPH, Alfa Aesar, Cat# 44150) radical scavenging assay[4]. Briefly, a 0.2 mM DPPH solution was prepared in 1:1 mixture of ethanol (Fischer Scientific, Cat# BP2818-500, 200 proof) and water (Fisher Scientific, Cat# W2-4) since SLS or TLS is not completely soluble in ethanol. SLS or TLS at 3 mg/mL was dissolved with $CaO_2$ at concentrations ranging from 0 to 0.5 mg/mL. DPPH solution without sample was used as the control. After incubating samples in darkness at room temperature for 30 min and 24 h with mild agitation of 150 rpm, decreases in absorbance were measured at 517 nm using Cytation3 spectrophotometer (Biotek, Winooski, VT, USA). *L*-ascorbic acid at 3 mg/mL (Sigma-Aldrich, Cat# A4403-100MG) was used as a positive control. Absorbance of all samples without DPPH was subtracted to correct the background absorbance at 517 nm. The DPPH radical scavenging activity (%) was calculated using the following formula: DPPH radical scavenging activity (%) = $(A_c - A_s)/A_c \times 100$ (%), where $A_c$ is absorbance of control and $A_s$ is absorbance of samples.

### SUPPLEMENTARY FIGURES

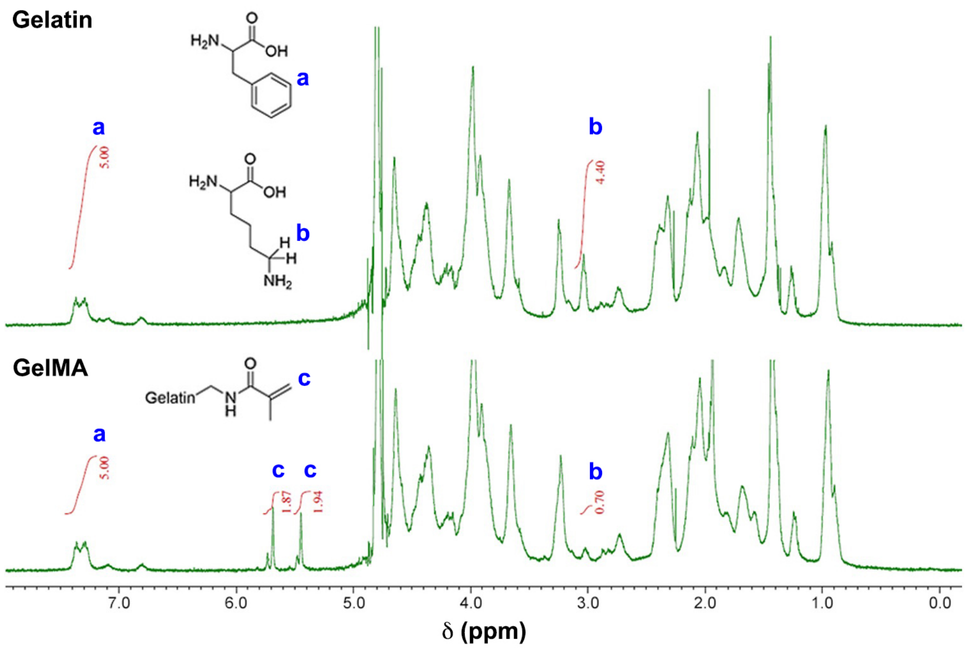


**Figure S1. $^1$H NMR ($D_2O$) spectrum of gelatin and GelMA.** Coupling of MA to gelatin was quantified from the integrated peak intensities of protons in phenyl group (a), alkene (b) and carbon adjacent to primary amine of lysine (c).

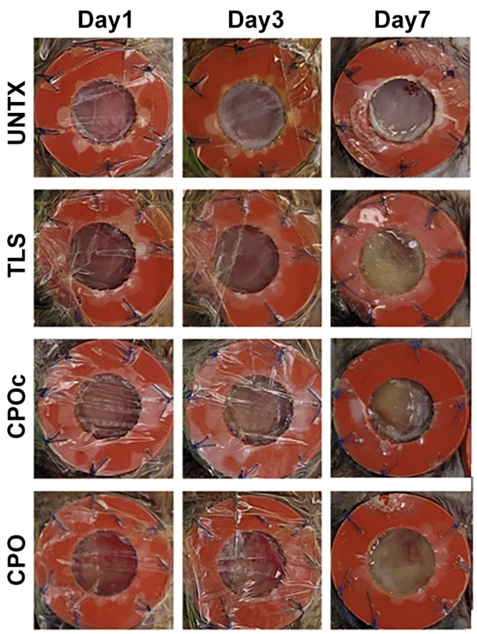


**Figure S2. Photographs of wounds after surgery.** Lignin composites were crosslinked *in situ* and wounds were monitored in wildtype C57BL/6N mice (8 to 10-week-old) up to 28 days. Photographs were taken at 1, 3 and 7 days after surgery. Details of compositions of UNTX, TLS, CPOc and CPO are in **Table 1**.

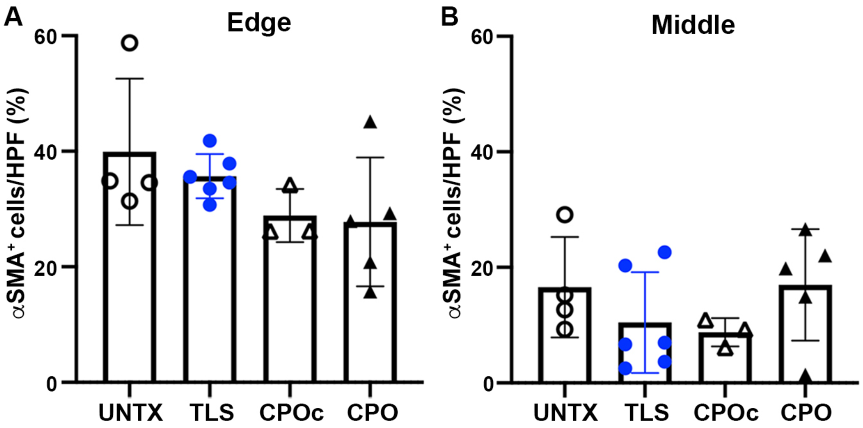


**Figure S3. Quantification of αSMA$^+$ cells in the wound treated with lignin composites.** αSMA$^+$ cells quantified in the edge sections (A) and middle sections (B) of the wound (total shown in **Figure 6**). Either one-way ANOVA with Tukey's HSD *post hoc* tests or with Kruskal-Wallis test followed by Dunn's test shows no statistically significant difference. $3 \leq n \leq 6$, mean±SD. Details of compositions of UNTX, TLS, CPOc and CPO are in **Table 1**.

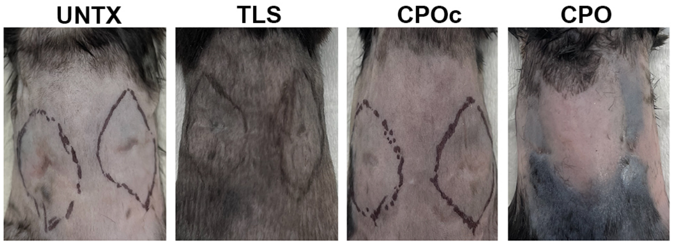


**Figure S4. Photographs of wounds at 28 days after surgery.** For scar assessment, photographs were taken from all 4 groups of treatment before harvest. Details of compositions of UNTX, TLS, CPOc and CPO are in **Table 1**.

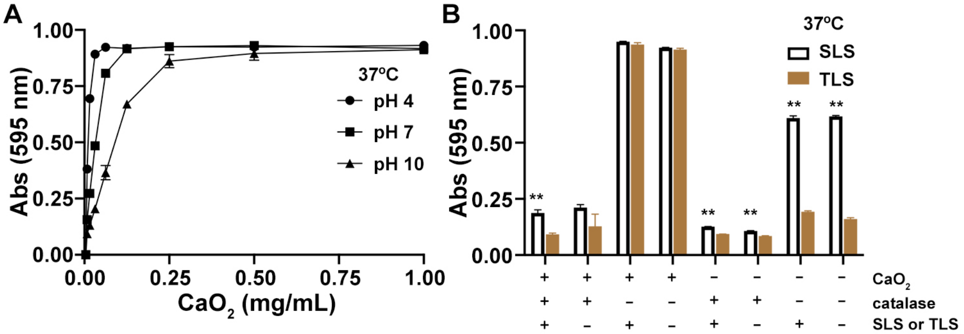


**Figure S5. Released peroxide from $CaO_2$ is dissociated into $O_2$ primarily by catalase and preferably at lower pH, which is further facilitated by TLS.** (A) The pH of PBS altered the released amount of peroxide from $CaO_2$. (B) The solubility limit of $CaO_2$ is 1.65 mg/mL. When $CaO_2$ was dissolved at PBS (pH 7.4, 37°C), the dissociation of peroxide to oxygen was catalyzed by catalase (100 U/mL). While neither SLS nor TLS significantly catalyzed the dissociation of peroxide to oxygen at 50 μg/mL of $CaO_2$, TLS facilitates the dissociation of peroxide to oxygen at 5 μg/mL of $CaO_2$. *First symbol*: +, $CaO_2$ 50 μg/mL; -, $CaO_2$ 5 μg/mL / *Second symbol*: +, catalase 100 U/mL; -, catalase 0 U/mL / *Third symbol*: +, SLS or TLS 3 mg/mL; -, SLS or TLS 0 mg/mL. Student's *t*-test, **$p<0.01$; n=3, mean±SD.

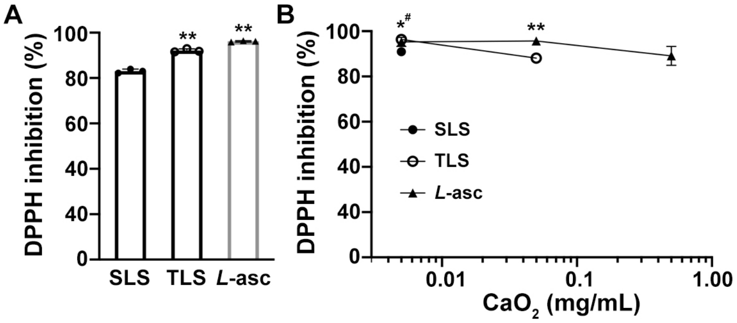


**Figure S6. Minor reduction of antioxidation capacity of SLS and TLS in the presence of $CaO_2$.** (A) DPPH inhibition in the absence of $CaO_2$, similar to the previously reported results[4]. (B) DPPH inhibition in the presence of $CaO_2$ at the concentrations ranging from 5 to 50 μg/mL. One-way ANOVA with Tukey's *post hoc* test, *$p<0.05$ and **$p<0.01$, #denotes the exception to the comparison of *L*-asc to TLS. n=3, mean±SD.

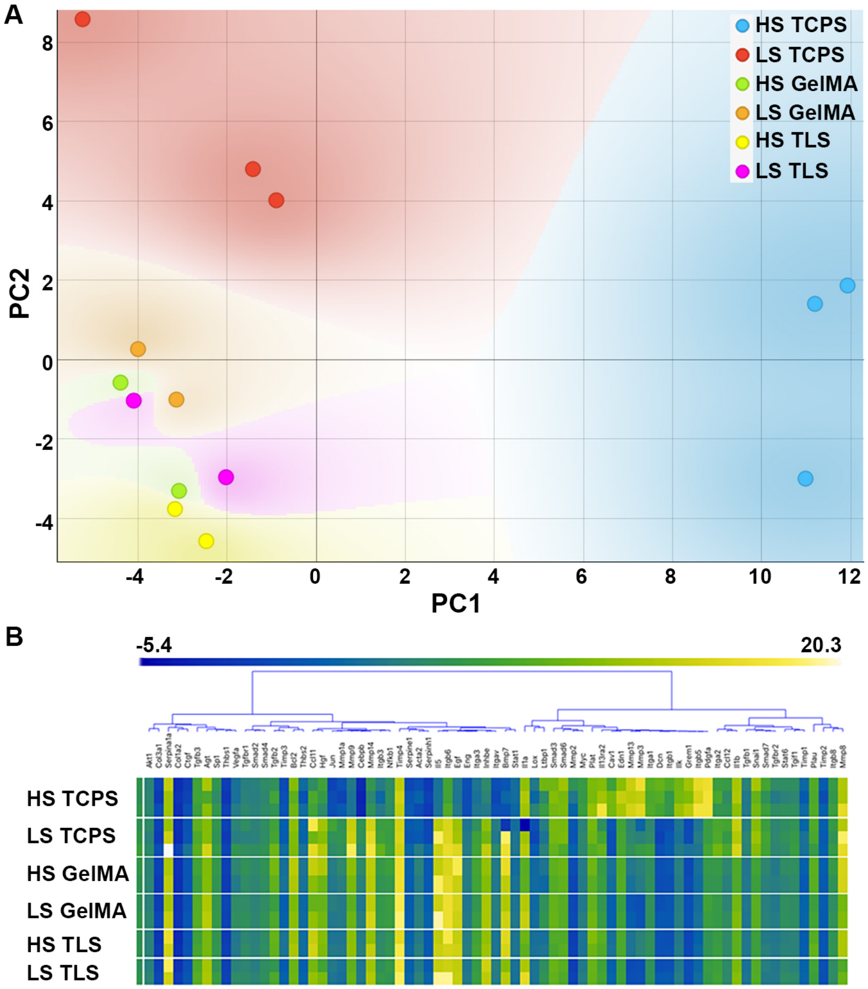


**Figure S7. Fibrosis PCR array of human dermal fibroblasts (hdFBs).** (A) Principal Component Analysis (PCA) of 84 fibrosis-related genes from high scar (HS) and low scar (LS) hdFBs cultured on three different substrates (TCPS, GelMA and TLS, see **Table 1**). (B) Hierarchical Clustering (HC) of 84 fibrosis-related gene. LS exhibits a proliferative phenotype. TLS lignin composites modulate the profiles of the fibrosis-related genes.
